# Cross-state stability and maturational similarity of the social brain connectome in children and adolescents

**DOI:** 10.64898/2026.10.13.758974

**Authors:** Jacqueline Lee, Jeff Eilbott, Clare Kelly, Rebecca Todd, Tamara Vanderwal

**Affiliations:** Department of Psychology, University of British Columbia, Vancouver, BC, Canada; Department of Psychiatry, University of British Columbia, Vancouver, BC, Canada; BC Children’s Hospital Research Institute, Vancouver, BC, Canada; School of Psychology, Trinity College Dublin, Dublin, Ireland; Department of Psychiatry, School of Medicine, Trinity College Dublin, Dublin, Ireland; Trinity College Institute of Neuroscience, Trinity College Dublin, Dublin, Ireland; Djavad Mowafaghian Centre for Brain Health, University of British Columbia, Vancouver, BC, Canada

**Keywords:** movie-fMRI, functional connectivity, whole brain, social brain, predictive modelling, development

## Abstract

The *intra-individual stability* of functional connectivity (FC) across two or more scans and the *inter-individual similarity* of FC patterns relative to peers capture developmental variability in functional brain organization. Despite their different referents, these measures are strongly coupled at the whole brain level; however, it remains unclear whether this relationship emerges in development, whether it extends to the social brain, and whether it differs by sex. Based on the importance of social development in this age range, we hypothesized that known developmental relationships and sex-specific effects would be more pronounced in the social brain relative to the whole brain.

Using a large psychiatrically enriched sample of 769 youth ages 6-21 (female N = 327, mean age = 12.27 years), we computed FC during rest and movie-watching scans and quantified stability across scans as well as similarity to mature subjects (n=36) within each condition. We then tested whether sex-dependent age relationships were captured by these measures or by the strength of their coupling.

Our results showed that cross-state stability and maturational similarity are strongly coupled in development, and that this coupling is enhanced in the social brain relative to the whole brain. Furthermore, the strength of stability-similarity coupling increased as a function of age in females, while males showed the opposite effect. When assessing relationships between age and each FC measure independently, no individual effects were significant after FDR correction. However, our multivariate age-prediction models fared much better, demonstrating that edgewise stability and maturational similarity measures can capture spatially heterogeneous developmental effects that are lost when collapsing across edges.

Together, these results support the emerging idea that the degree of FC stability across scans is related to how closely an individual’s functional architecture aligns to a shared template. Here, we newly show that this stability-similarity coupling is stronger in the social brain and varies with age in sex-specific ways. Future longitudinal work is needed to establish whether these cross-sectional relationships also characterize within-person developmental change.

## 1. Introduction

### 1.1 Probing functional brain development using intra- and inter-subject measures

For decades, researchers have known that test-retest reliability of resting state connectomes in children and youth is lower than in adults (Somandepalli et al. 2015). Although this was initially attributed to noise, more recent work suggests that this variability in pediatric functional connectivity (FC) reflects developmentally meaningful differences in *intra-subject stability*, measured as the correlation between a subject’s FC across scans, and *inter-subject similarity*, measured as the correlation between a subject’s FC to their peers within a scan.

Past work has primarily assessed these measures at the whole brain level. Developmental studies have examined intra-subject stability and inter-subject similarity across repeated resting state scans, across tasks, and across movies, and found that these measures relate to age, symptoms and/or behaviors (Kaufmann et al., 2017; Vanderwal et al., 2021; Fu et al., 2023; Horien et al., 2019; Graff et al., 2022a). To begin, Kaufmann et al. (2017) found that the average connectome stability across different tasks was positively associated with age and negatively associated with psychopathology in youth ages 8-22 (N=797). In our earlier work with a smaller sample (Vanderwal et al., 2021), we found that connectome stability across rest scans and connectome similarity during movie-watching were positively associated with social abilities but not age or psychopathology in a help-seeking sample of youth ages 6-21 (N=210). More recently, Hafeman et al. (2025) found that longitudinal stability within networks-of-interest distinguished youth with bipolar disorder from typically developing participants with excellent accuracy across rest scans. Together, these works suggest that connectome stability within a subject and connectome similarity between subjects relate to brain function in various ways.

Two methodological choices are pivotal when interpreting these measures or comparing findings across studies. First, acquisition state changes the central meaning of the term stability. Intra-subject stability assessed across the same scan type (e.g. rest-to-rest) indexes the reliability of an individual’s FC pattern within a given state, whereas stability across different scan types (e.g. rest-to-movie) indexes how strongly that pattern is reorganized across states. In contrast, inter-subject similarity is almost always assessed within a given scan. During rest, similarity may reflect shared aspects of intrinsic network architecture, whereas during task, similarity may also capture shared task- or stimulus-driven FC organization. Second, the interpretation of inter-subject similarity depends on the reference group to which each individual is compared. In developmental FC studies, similarity is often measured relative to the group as a whole, indexing the “typicality” of an individual’s functional architecture (Kaufmann et al., 2017; Vanderwal et al., 2021; Graff et al., 2022a). In developmental inter-subject correlation (ISC) studies, time-course similarity has been quantified relative to mature subjects to capture “neural maturity” (Cantlon and Li, 2013; Richardson et al., 2018). Here, we extend this maturity approach to FC, providing a new analytical framework to assess *maturational similarity*, which may yield novel insights regarding how and when FC patterns converge towards a common template.

### 1.2 Relationships between intra-subject stability and inter-subject similarity

Relationships between FC stability and FC similarity have been indirectly examined using connectome fingerprinting, which asks whether individuals can be correctly identified from the group across scans based on their unique patterns of FC (Finn and Shen et al., 2015). Successful identification depends on the balance between intra-subject stability and inter-subject similarity such that an individual’s FC must be more similar to their own pattern across scans than to the patterns of other participants. Despite developmental variability of FC, whole brain connectomes have been shown to be highly identifiable in youth, even when scans are separated by months or years (Jalbrzikowski et al., 2020; Horien et al., 2019; Demeter et al., 2020; Camp et al., 2024). These findings indicate that functional organization remains uniquely individualized during periods of substantial developmental change.

At the subject level, individualization has also been quantified by taking the difference or the ratio between an individual’s stability and group similarity (e.g. Corriveau et al., 2023; Graff et al., 2022a). For instance, Graff et al. (2022a) found that across two movie-watching scans collected 12-months apart, typically developing children 4-7 years old (N=73) had 98% fingerprinting accuracy, and that stability and individualization (stability-similarity difference) were positively associated with age. This identifiability is also reflected in the finding that intra-subject stability tends to be higher than inter-subject similarity in youth (e.g. Graff et al., 2022a; Hafeman et al., 2025).

Other recent work has directly examined relationships between intra-subject stability and inter-subject similarity by testing for correlations between these measures. Somewhat surprisingly, FC stability and FC similarity show strong positive correlations within individuals (Vanderwal et al., 2021; Liao et al., 2021; Corriveau et al., 2023), suggesting that people with more stable connectivity profiles also tend to exhibit more group-typical patterns. However, no studies to date have explored whether FC stability and maturational similarity are coupled in development or whether this relationship extends to social brain systems, which undergo major refinement in childhood and adolescence (Andrews et al., 2021).

### 1.3 The social brain during childhood and adolescence

The ‘social brain’ is a distributed network of regions involved in an array of social and affective processes. Nodes of this network include but are not limited to the temporoparietal junction (TPJ), medial prefrontal cortex (mPFC), precuneus, posterior superior temporal sulcus (pSTS) and inferior frontal gyrus (IFG) (Blakemore, 2008). An unexpectedly low number of studies have probed social brain development using FC, particularly across a full social brain connectome that includes both higher- and lower-level systems (e.g., fusiform gyrus, middle temporal V5 area; Alcalá-López et al., 2018). To our knowledge, only two studies have assessed FC within a “complete” social brain network during development (McCormick et al., 2018; Saljoughi et al., 2026), and both used standard FC approaches to assess inter-regional connectivity. However, these studies did not investigate the functional maturity of social brain FC or the stability of these patterns across different states.

Movies can act as ‘amplifiers’ of individual differences in FC (e.g. Vanderwal et al., 2017; Finn and Bandettini, 2021) and have been shown to capture developmental differences that are less apparent during rest or more constrained task paradigms (e.g. Cantlon and Li, 2013; Richardson et al., 2019). Movies are also potent drivers of social processing and have been used in many developmental studies (Redcay and Moraczewski, 2020). For example, several movie-fMRI studies have demonstrated that higher-level social brain regions (e.g., the TPJ and mPFC) show increasing levels of functional specialization and more mature neural responses with age and social cognitive abilities (Saxe et al., 2009; Gweon et al., 2012; Richardson et al., 2018, 2019; Moraczewski et al., 2018, 2020). However, no work to date has assessed whether these developmental relationships are also captured by functional maturity of social brain FC patterns.

Finally, while sex-based differences in social brain functioning have been well characterized in adults (Proverbio, 2023), few studies have assessed this in developmental samples (although see Mills et al., 2014 for structural findings). Given that social functioning has been shown to differ between males and females across development (Calero et al., 2013; Poznyak et al., 2019; Desatnik et al., 2023), characterizing sex-dependent developmental relationships in social brain functioning may provide novel insight into these differences at the neurobiological level.

### 1.4 Study Aims

The goal of this current work is to provide the first assessment of FC stability and FC maturational similarity within the social brain, to compare these developmental relationships to whole brain patterns, and to assess sex-specific effects. We use a social brain atlas that contains regions across the full hierarchy of social processing (i.e., both lower-level and higher-order regions) and apply it to a large sample (N=769) of psychiatrically enriched youth, ages 6-21 years from the Healthy Brain Network (HBN) Biobank (Alexander et al., 2017).

First, we assess whether social brain FC is individualized across rest and movie scans using binary (fingerprinting) and continuous (discriminability) measures of identifiability.

Although the social brain connectome comprises only 36 regions, it includes some of the most densely connected regions in the brain, spanning multiple levels of the cortical processing hierarchy (Alcalá-López et al., 2018). It also overlaps with frontoparietal regions which have been repeatedly shown to enable successful fingerprinting in adults (Finn & Shen et al., 2015; Vanderwal et al., 2017). Thus, we hypothesized that this distributed architecture would capture individualized patterns of connectivity across states, even though it contains fewer regions.

Next, per subject, we compute for whole brain and social brain: one measure of *cross-state stability* to index FC modulation across rest and movies, and two measures of *within-state similarity* to index FC maturational similarity during rest and movies. To our knowledge, this is the first study to apply this “similarity to peers” approach using social brain FC. In our main analyses, we test whether developmental variability is captured by cross-state stability, maturational similarity, or the strength of their coupling, and examine whether these relationships show sex-specific effects. As a secondary outcome, we also measure total t-scores from the Social Responsiveness Scale-2 (SRS; Constantino et al., 2003) and examine whether individual differences in social functioning relate to stability and maturational similarity.

We predicted that maturational similarity would increase with age and social functioning, informed by previous developmental ISC work (e.g. Richardson et al., 2018). We also anticipated that these associations would be more pronounced during movie-watching compared to rest, consistent with findings in which movies enhanced sensitivity to individual differences (e.g. Finn and Bandettini, 2021). With regards to cross-state stability, we predicted that higher stability would be associated with higher age and social functioning based on previous developmental work assessing FC stability across tasks (Kaufmann et al., 2017), repeated movie scans (Graff et al., 2022a), and repeated rest scans (Vanderwal et al., 2021). Given the importance of social development across this age range, we also anticipated that the social brain would capture partially distinct and potentially stronger developmental effects compared to the whole brain. We also predicted that social brain FC measures would track differently with age as a function of sex, consistent with behavioural studies on social development.

As a final exploratory aim, we derive edgewise measures of stability and maturational similarity and use multivariate prediction models to test whether they predict age. We anticipated that these edgewise FC measures would significantly predict age and that developmental relationships would show spatially heterogeneous effects across the brain.

## 2. Methods

### 2.1 Methods Overview

Using FC derived during rest and movie-watching scans, we first assess connectome individualization using fingerprinting and discriminability. For each individual and connectome type (whole brain and social brain), we calculate one measure of *cross-state stability* by correlating each subject’s rest and movie FC, and two measures of *within-state similarity* by correlating FC from each subject to the mature subgroup for rest and movies separately. Using these summary FC measures, we perform linear-mixed effects (LME) models to assess associations between stability and similarity during development and whether the strength of this stability-similarity coupling is related to age as a function of sex. We then use LME models to assess sex-stratified associations with age for each stability and similarity measure independently, as well as associations with social functioning scores. Finally, we perform multivariate predictive modelling of age using whole brain and social brain edgewise stability and similarity measures. Using our whole brain models, we assess the spatial distribution of highly predictive edges and their correspondence to our social brain mask.

### 2.2 Sample and Data

Data were obtained from the Healthy Brain Network (HBN), a publicly available dataset containing a large psychiatric help-seeking sample of youth aged 6–21 years recruited from the greater New York City area (Alexander et al., 2017). Study procedures were approved by the Chesapeake Institutional Review Board. Written informed consent was obtained from participants aged 18 years and older, otherwise, parental consent with participant assent was obtained.

Using HBN releases 1–11, functional MRI data across 4 runs were obtained in a fixed order: two sequential 5.1-minute (375 volumes) eyes-open rest scans, a 10-minute (750 volumes) movie-watching scan during which a clip from *Despicable Me* was shown, and a 3.5-minute movie-watching scan (250 volumes) during which a short film called *The Present* was shown.

Our previous study utilized all 4 fMRI runs allowing for estimation of rest-to-rest and movie-to-movie stability (Vanderwal et al., 2021). Here, we chose to omit these pairings due to reliability concerns as they relied on 5 minutes or less of data (Noble et al., 2017). After discarding the first five volumes from the two rest and one movie run, the two rest runs were concatenated, resulting in a single rest run with 740 volumes, and a movie-watching run with 745 volumes.

A total of 923 participants with complete resting-state and movie-watching data were initially identified. Participants were excluded if either run met one or more of the following criteria: (i) mean root-mean-squared displacement (RMSD; Jenkinson et al., 2002) greater than 0.25 mm; (ii) fewer than 7 minutes (525 volumes) of usable data remaining after motion censoring using an RMSD threshold of 0.25 mm and removing the affected volume and one before and after; or (iii) poor spatial coverage, defined as missing nodes for any whole brain parcel or less than 50% coverage for any social brain ROI. An additional exclusion was applied for missing basic demographic information (age, sex, or site).

The final sample consisted of 769 participants. Phenotypic data were collected in Staten Island (n=251), Midtown (n=263), or Harlem (n=255). MRI scans were collected at Rutgers University Brain Imaging Center (RUBIC; *n* = 254), Cornell Brain Imaging Center (CBIC; *n* = 450), or City University of New York (CUNY; *n* = 65).

### 2.3 Behavioral Measures

Total t-scores from the Social Responsiveness Scale-2 (SRS; Constantino et al., 2003) were used to assess social functioning for subjects with valid data (N=736). Although originally developed to quantify social deficits associated with Autism Spectrum Disorder, its use as a dimensional measure of social functioning in both clinical and non-clinical populations is supported (Bruni, 2014; Hus et al., 2013). Additionally, overall levels of psychopathology were measured using total t-scores from the Child Behavioural Checklist (CBCL; Achenbach, 1999) – CBCL scores were only used to identify subjects below the clinical cut-off within the mature subsample.

### 2.4 MRI Acquisition and Preprocessing

MRI data were collected either using a Siemens 3 Tesla Tim Trio Scanner or a Siemens 3 Tesla Prisma scanner. A 3D T1-weighted structural MP-RAGE volume was acquired with TR = 2500 ms, TE = 3.15 ms, FA = 8°, slice thickness = 0.8 mm, slices = 224. Echo-planar imaging sequences for all functional runs were acquired with TR = 800 ms, TE = 30 ms, FA = 31°, slice thickness = 2.4 mm, number of slices = 60, FOV = 204 mm, multiband factor = 6, and voxel size = 2.4 x 2.4 x 2.4 mm.

Preprocessing was performed using fMRIPrep version 22.1.1 (Esteban et al., 2019) including slice time correction, motion realignment, and transformation into Montreal Neurological Institute space (2 mm MNI template). Denoising was performed using 3dTproject in Afni version 25.1.10 (Cox, 1996) by simultaneously regressing out all confound parameters, which included the 24 head motion parameters (6 motion parameters, their derivatives and squared derivatives), the first 10 anatomical CompCor components, the global signal, and constant, linear and quadratic trends. Temporal filtering was implemented in the same regression model using band-pass filter basis functions (0.01 and 0.10 Hz). To further mitigate the impact of motion in this clinical, developmental sample, we performed moderate motion censoring using an RMSD threshold of 0.25 mm, removing the volume before and after. We elected to use global signal regression (GSR) to align with previous work demonstrating that censoring, head motion parameter regression, bandpass filtering, and GSR provides the best balance of noise removal and information recovery in pediatric samples (Graff et al., 2022b).

### 2.5 Generating Whole Brain and Social Brain Connectomes

For parcellation at the whole brain level, we used a 268-node atlas (Shen et al., 2013) that covers cortical, subcortical, and cerebellar regions. Nodes were assigned to 10 large-scale functional networks as previously done (e.g. Noble et al., 2017). For parcellation at the social brain level, we used a 36-region atlas (Alcalá-López et al., 2018) derived from task-activation convergence from 3,972 neuroimaging studies across a broad range of social-affective paradigms. Four social brain sub-networks (visual-sensory, limbic, intermediate, and higher-level associative networks) were derived based on similar patterns of connectivity across tasks and resting-state using a hierarchical clustering analysis.

Whole brain (268 x 268 FC) and social brain (36 x 36) FC matrices were constructed for each participant and condition (Rest and Movie) by averaging voxel timeseries within each node/region and then correlating across all node/region pairs using Pearson’s r. Correlation coefficients were Fisher z-transformed, and the values from the lower triangle of the FC matrices were vectorized, yielding 35,778 unique edges for whole brain FC and 630 unique edges for social brain FC. These z-transformed vectors were carried forward for all subsequent FC analyses (see *Figure 1A*). This approach to the social brain is quite restrictive: FC for a given region reflects its connectivity to *only* regions within the social brain, and it does not capture connections between those regions and the rest of the brain as is often done (e.g., when masking for a dual-regression analysis).

**Figure 1.**
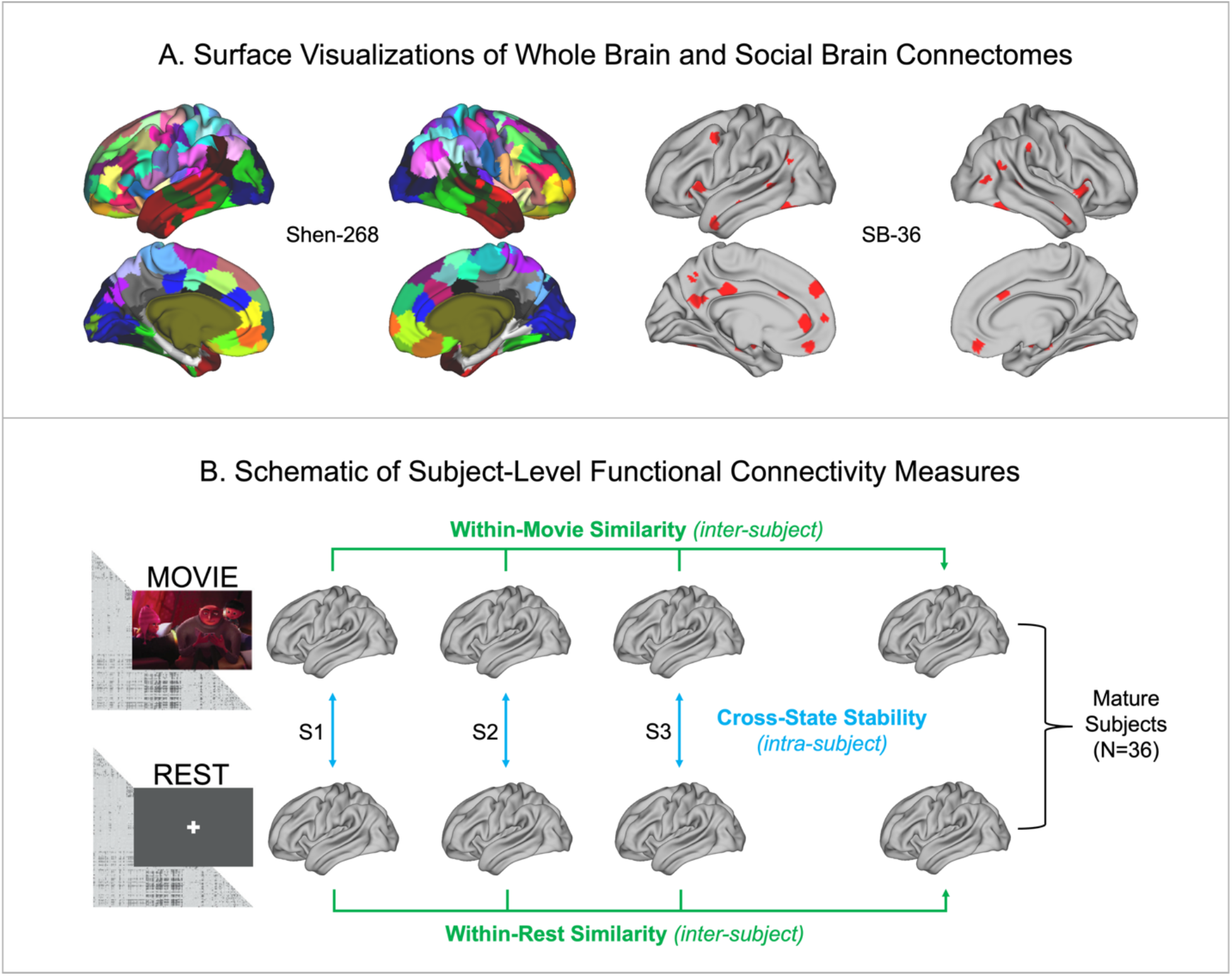
Connectome Surface Visualizations and Functional Connectivity Metrics. **(A)** Surface visualizations of the whole brain (Shen-268) and social brain (SB-36) connectomes. The Shen-268 and SB-36 atlases are displayed on fs_LR 32k midthickness surfaces using Connectome Workbench. **(B)** Cross-state stability was calculated as intra-subject similarity of functional connectivity patterns across movie and rest scans. Within-state similarity measures were calculated as inter-subject similarity of functional connectivity patterns to subjects in the healthy mature reference group (n=36) during movie viewing (top row) and rest (bottom row).

### 2.6 Measuring Connectome Individualization Across States

Connectome individualization was assessed using fingerprinting and discriminability measures. Both are non-parametric, multivariate summary metrics used to evaluate the extent to which each person’s connectome is distinct from others in the group. Fingerprinting provides a binary test of identifiability; i.e., it asks whether a participant’s connectome is most similar to their own data from another run, while discriminability measures identifiability on a continuous scale, i.e., it asks how much more a participant resembles themself compared to others across runs. For both measures, identifiability was assessed across states with each scan serving as ‘test’ and ‘target’ databases. To assess significance, nonparametric permutation testing was performed by randomly shuffling participant identity labels in the target database and recomputing fingerprinting and discriminability measures. We performed 1000 permutations to generate null distributions and obtained p-values as the proportion of permuted values greater than or equal to the observed individualization measurement.

#### 2.6.1 Fingerprinting

Fingerprinting was performed following procedures described by Finn & Shen et al. (2015). For each participant’s connectome in the ‘test’ database, similarity to all participants’ connectomes in the ‘target’ database was calculated between pairs of FC vectors using Pearson’s correlation. A participant was considered successfully identified if the highest correlation was obtained with their own connectome in the target database. Fingerprinting accuracies were calculated *across states* as the proportion of successful identifications across all participants. To assess whether head motion might contribute to successful matching, identification accuracies were also computed using the distribution of RMSD values (in place of FC values).

#### 2.6.2 Discriminability

Discriminability analyses were performed closely following procedures by Bridgeford et al. (2021), however, rather than using Euclidean distance, we used similarity (i.e. Pearson’s correlation) as our unit of measurement. First, correlations between a participant’s connectome in the database and all connectomes in the target set were computed, resulting in one within-participant and N–1 between-participant similarity values for each participant. Discriminability was then calculated as the number of observations where between-participant similarity was less than within-participant similarity over the total number of comparisons.

#### 2.6.3 Differential Power

To assess edges with the highest discriminability, we computed the differential power (DP) of each edge following previously used methods (Finn & Shen et al., 2015, Vanderwal 2017; Horien et al., 2019). First, FC vectors were normalized within each participant and condition. Then for each edge, an N x N participant similarity matrix was constructed to capture within-subject similarity and between-subject similarity values. At the subject-level, DP for each edge was calculated as the proportion of between-participant similarities that exceeds within-participant similarity. These DP values were -log transformed and summed across participants to estimate DP of each edge at the group-level. Edges with higher DP values are interpreted as contributing more strongly to participant identification by consistently distinguishing individuals from one another. Edges were ranked according to DP magnitude and edges in the 95^th^ percentile were selected for subsequent visualization of network-level distributions.

### 2.7 Cross-State Stability & Within-State Similarity

For each participant and connectome type (whole brain and social brain), we computed one measure of cross-state stability and two measures of within-state similarity relative to mature subjects (for rest and movie). Cross-state stability was calculated as the Pearson correlation between a participant’s rest and movie connectomes. Within-state similarity for each condition was calculated as the average Pearson correlation between each participant’s connectome to all other individuals in the mature subsample. Within-state similarity for participants in the mature subsample was calculated using pairwise correlations to all other mature subjects, excluding oneself. The mature subsample included participants 17 years and older (N=36) with scores below clinical cut-offs for SRS (< 60) and CBCL (< 65). See *Figure 1B* for a schematic.

### 2.8 Stability and Similarity Relationships

To examine relationships between stability and similarity measures, we used a LME model with Restricted Maximum Likelihood (REML) using fitlme in MATLAB. We modelled cross-state stability as a function of mean within-state similarity (averaged across rest and movie), with connectome type (whole brain vs social brain) as an interaction term, age, sex, and RMSD as covariates, and site and participant as random intercept factors. All continuous variables were z-scored prior to modelling. Fixed effects and model-derived linear contrasts were used to assess similarity–stability relationships for whole brain and social brain, as well as differences between connectome types. In all iterations of the LME model, whole brain was set as the reference level. When applicable, model-based simple slope tests were performed using coefTest in MATLAB to evaluate conditional similarity–stability slopes. To assess whether the strength of stability-similarity relationships vary as a function of age and sex, we repeated the LME models using age as a moderator for the full-sample and stratified by sex.

### 2.9 Age and SRS Associations

Again using LME models with REML, we modeled each FC measure as a function of age or SRS scores, with connectome type as an interaction term, age (only for SRS models), sex, and RMSD as covariates, and site and participant as random intercept factors. Prior to modelling, SRS scores were flipped so that higher scores indicated better social functioning. To explore sex-dependent developmental effects, we also assessed sex-stratified relationships. All continuous variables were z-scored prior to modeling. Models were fit separately for each FC measure and each phenotypic measure of interest. Furthermore, due to the tight coupling between stability and similarity measures, we controlled for average within-state similarity in models assessing cross-state stability and vice versa for our within-state similarity models (as done in prior work, see Liao et al., 2021). To account for multiple comparisons, *p*-values from the LME models were adjusted using the Benjamini–Hochberg false discovery rate (FDR) procedure (Benjamini and Hochberg, 1995). Results were considered statistically significant at an FDR-corrected threshold of *q* < 0.05.

### 2.10 Multivariate Predictions of Age

Summary measures of stability and similarity collapse information across all edges and do not account for spatially heterogeneous effects. In other words, two participants may have the same average level of stability or similarity, but differ in which specific connections show stronger or weaker stability/similarity. We therefore sought to determine whether retaining edgewise information could better capture complex and spatially distributed developmental patterns. To do this, we derived vectorized measures of stability and similarity and used these edgewise measures to predict age. This approach allowed us to test whether developmental variation is encoded in multivariate spatial patterns of FC stability and similarity.

All edgewise FC measures were derived using distance-based similarity. Cross-state stability was calculated for each edge as the negative absolute difference between a participant’s movie and rest FC values. Within-state similarity (for movie and rest) was calculated for each edge as the mean negative absolute difference between a participant’s FC value and that of all participants in the healthy mature reference subgroup. These edgewise stability and similarity vectors were then used to predict age with regularized ridge regression using glmnet in R v4.5.2 (Friedman et al., 2010). Our procedures closely followed those described in previous studies (e.g. Adkinson et al., 2024; Gao et al., 2019). All models were evaluated in repeated 10-fold cross-validation. Within each cross-validation fold, we performed univariate feature selection (i.e., the processes of selecting which edges are retained in the model) using only the training participants. For each edge, we computed the partial Pearson correlation between the edge value and age, controlling for motion (mean RMSD). Edges were retained if their partial-correlation *p*-value was below α = 0.05. Within each fold, ridge regression models (*α* = 0 in glmnet) were then fit on the selected training edges. To determine the degree of penalization, ridge regression models were tested with 100 values of the penalty parameter λ (chosen by glmnet’s algorithm). The λ value that minimized the mean-squared error across all folds was selected as the best performing λ and each fold was then refit using this final λ to generate predictions for the held-out test fold. This procedure was repeated for 100 repetitions with differing fold assignments.

To assess prediction accuracy, out-of-fold predictions were evaluated against true age values using Pearson correlation (*r*) and root-mean-squared error (RMSE). Statistical significance was assessed using non-parametric permutation testing. We performed 1000 label permutations where age values were randomly shuffled across participants. Ridge regression models were refit and prediction accuracy measures were recomputed for each permutation to generate null distributions. *P*-values were calculated as the proportion of the permuted outcomes that were greater (for r) or less than (for RMSE) the true mean prediction accuracy (averaged across the 100 model repetitions). To assess whether predictive performance differed between within-rest and within-movie similarity models, we performed paired t-tests to compare r and RMSE values from the 100 model repetitions.

### 2.11 Data and Code Availability

Data used in this study are publicly available through the HBN scientific data portal (https://fcon_1000.projects.nitrc.org/indi/cmi_healthy_brain_network/). Participant IDs and code for the analyses can be found on the following GitHub repository: https://github.com/jac-lee-ubc/fc-stability-similarity. Parcellations and node-network assignments for the Shen whole brain atlas were obtained on NITRC and GitHub (https://www.nitrc.org/frs/?group_id=51; https://github.com/SNeuroble/atlas_tools/tree/master). The social brain ROIs from the Alcalá-López et al (2018) social brain atlas were obtained on NeuroVault (https://neurovault.org/collections/2462/).

## 3. Results

### 3.1 Participant Characteristics

Our sample contained 769 participants (327 female), aged 6–21 years (mean age = 12.27 ± 3.51 years). Paired t-tests demonstrated that mean RMSD for rest (0.10 ± 0.04 mm) and movie (0.10 ± 0.04 mm) were comparable across our final sample (*t*(768) = 1.53, *p*=0.127). Mean RMSD (averaged across rest and movie) was 0.10 ± 0.03 mm. As typical with pediatric samples, head motion (mean RMSD) was negatively correlated with age (*r* = -0.38, *p* = 2.68 x 10^-28^). Age distribution and head-motion characteristics are shown in *Figure 2*. SRS scores were not correlated with head motion (*r* = 0.064, *p*=0.08) or with age (*r* = 0.010, *p* = 0.79), see *Supplementary Figure 2* for SRS distribution and head-motion relationships.

**Figure 2.**
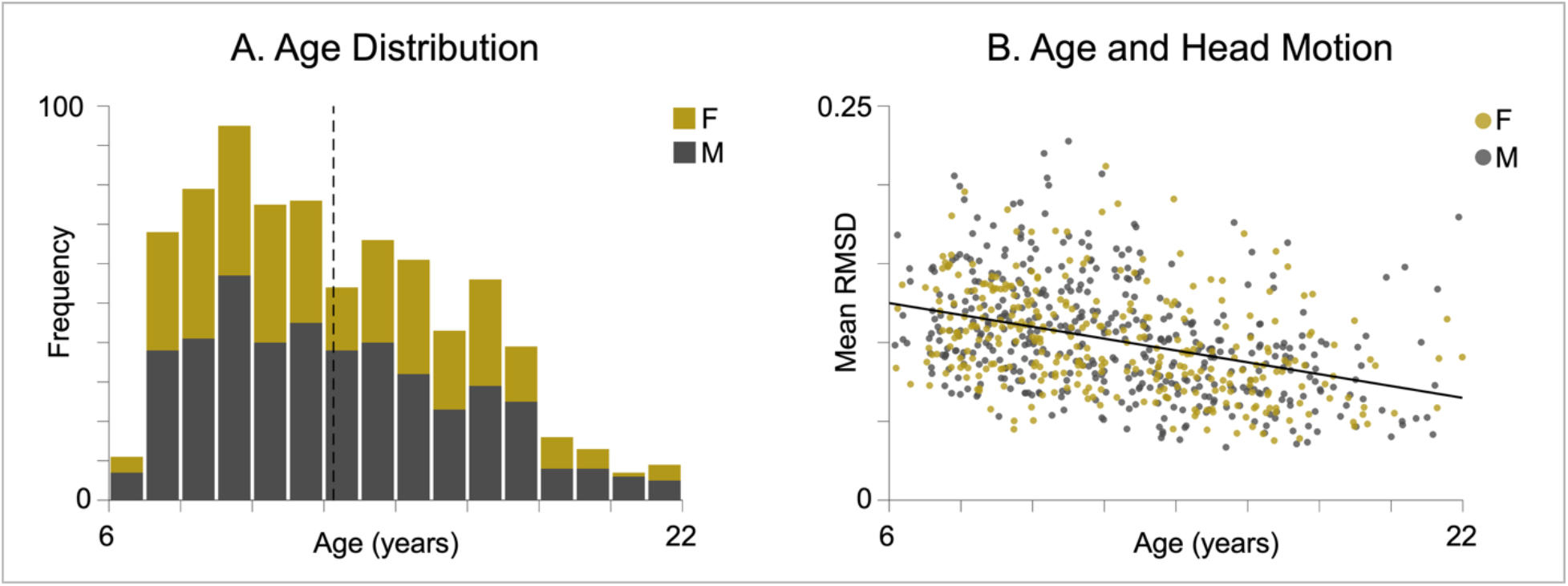
Sample demographics of the 769 participants (327 female). **(A)** Age distribution is visualized by sex; Mean age = 12.27 ± 3.51 (dashed line). **(B)** Relationship between age and head motion estimated using root-mean-squared displacement (RMSD). As expected, there was a strong negative correlation between age and mean RMSD (*r* = -0.38, *p* = 2.68 x 10^-28^).

Consistent with the HBN’s recruitment strategy, our sample was psychiatrically enriched, with most participants meeting criteria for one or more psychiatric diagnoses; the most common being Attention-Deficit/Hyperactivity Disorder, Inattentive Type (see *Supplemental Figure 1*).

The mature subsample consisted of 36 participants (13 female) from 17.01–21.56 years (mean age = 18.30 ± 1.40 years). All subjects in the mature subsample had SRS and CBCL total t-scores below clinical thresholds (mean SRS = 48.33 ± 5.61; mean CBCL = 49.64 ± 10.35).

### 3.2 Cross-State Connectome Individualization

Cross-state fingerprinting accuracies were all significantly higher than chance (*p*<0.001), ranging from 73.2-80.6% for the whole brain and 37.2-40.8% for the social brain. Discriminability values were all significantly higher than chance (*p*<0.001), ranging from 0.991-0.997 for whole brain and 0.958-0.970 for social brain. Head motion distribution vectors did not result in successful identification (2.86-3.38%). See *Figures 3A/B*.

**Figure 3.**
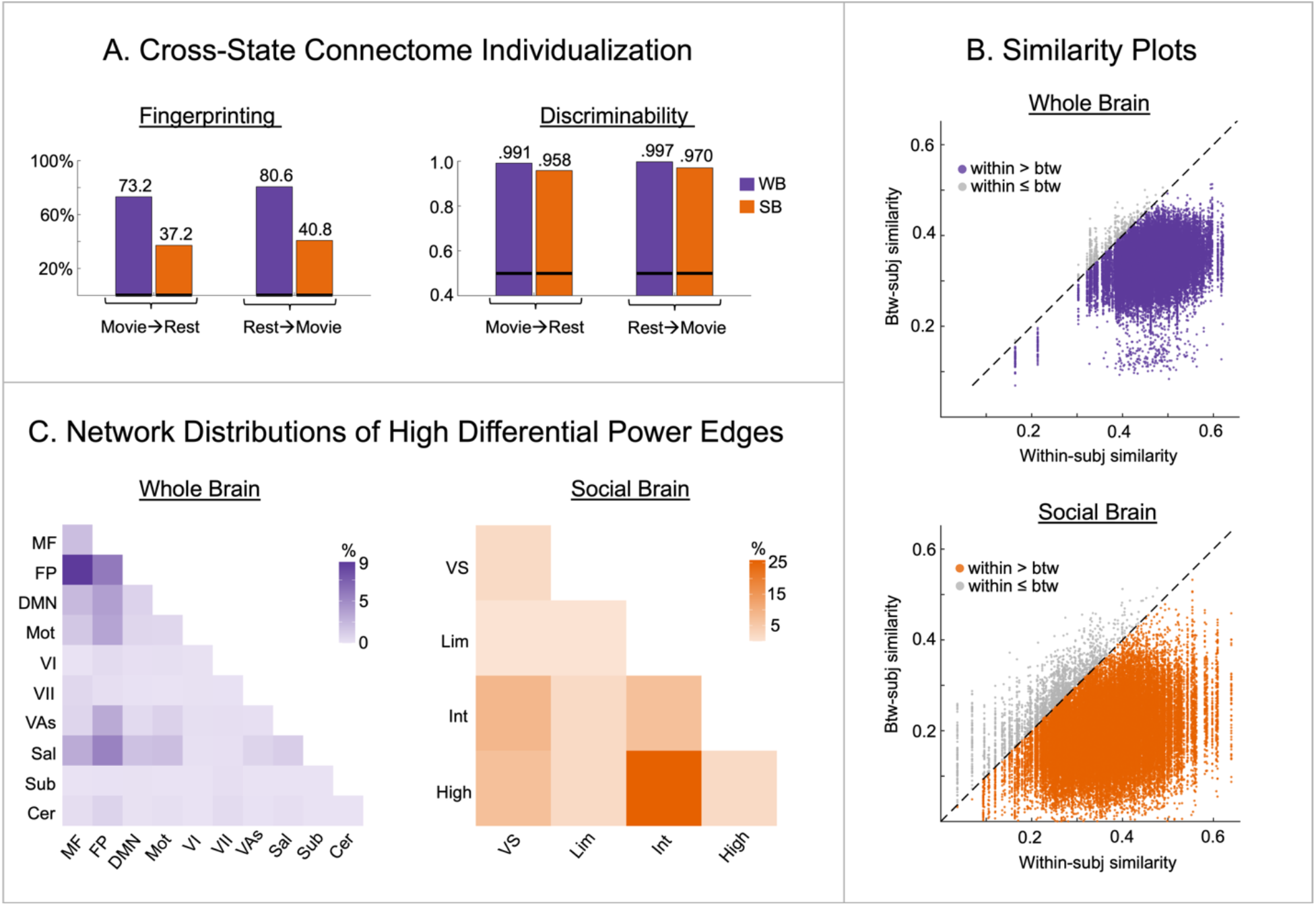
Connectome Individualization across Rest and Movie. **(A)** Fingerprinting accuracy and discriminability for whole brain (WB; 268 parcels) and social brain (SB; 36 parcels) connectomes. Black horizontal lines denote chance level accuracies and discriminability values obtained with permutation testing. **(B)** Two-dimensional similarity plots for whole brain (top) and social brain (bottom) connectomes (Movie = database, Rest = target). Each point represents the relationship between a subject’s within-individual similarity (x-axis) and between-individual similarity (y-axis). Points below the diagonal indicate instances where within-individual similarity exceeds between-individual similarity. Group discriminability = proportion of points below the diagonal; Fingerprinting accuracy = proportion of subjects whose entire column lies below the diagonal. **(C)** Network-level distribution of edges in the 95^th^ percentile of high differential power (DP) edges. Whole brain networks: MF, medial frontal; FP, frontoparietal; DMN, default mode; Mot, Motor; VI, visual I; VII, visual II; VAs, visual association; Sal, salience; Sub, subcortical; Cer, cerebellar. Social brain networks: VS, visual-sensory; Lim, limbic; Int, intermediate; High, higher-level.

Network distribution of edges with high DP were visualized in the main text using a 95^th^ percentile threshold (See *Figure 3C*). Whole brain discriminatory edges were mainly distributed across higher-level associative regions, particularly between frontoparietal and medial-frontal networks. At the social brain level, highly discriminatory edges were concentrated between high- and intermediate-level networks. See *Supplementary Figure 3* for DP results when thresholding at 90^th^ and 99^th^ percentiles.

Together, these findings demonstrate that whole brain and social brain connectomes show high discriminability across rest and movie states in youth ages 6-21, particularly across higher-level association regions. In contrast to the whole brain, fingerprinting accuracies were much lower using social brain FC, suggesting that continuous measures of identifiability better capture uniqueness at the social brain level.

### 3.3 Stability and Similarity Relationships

Prior to assessing stability-similarity relationships, we compared distribution patterns of each FC measure between the whole brain and social brain using paired t-tests and Pitman-Morgan tests. We found that group means, particularly for our stability measure, were lower and more variable across subjects in the social brain. See *Figure 4*.

**Figure 4.**
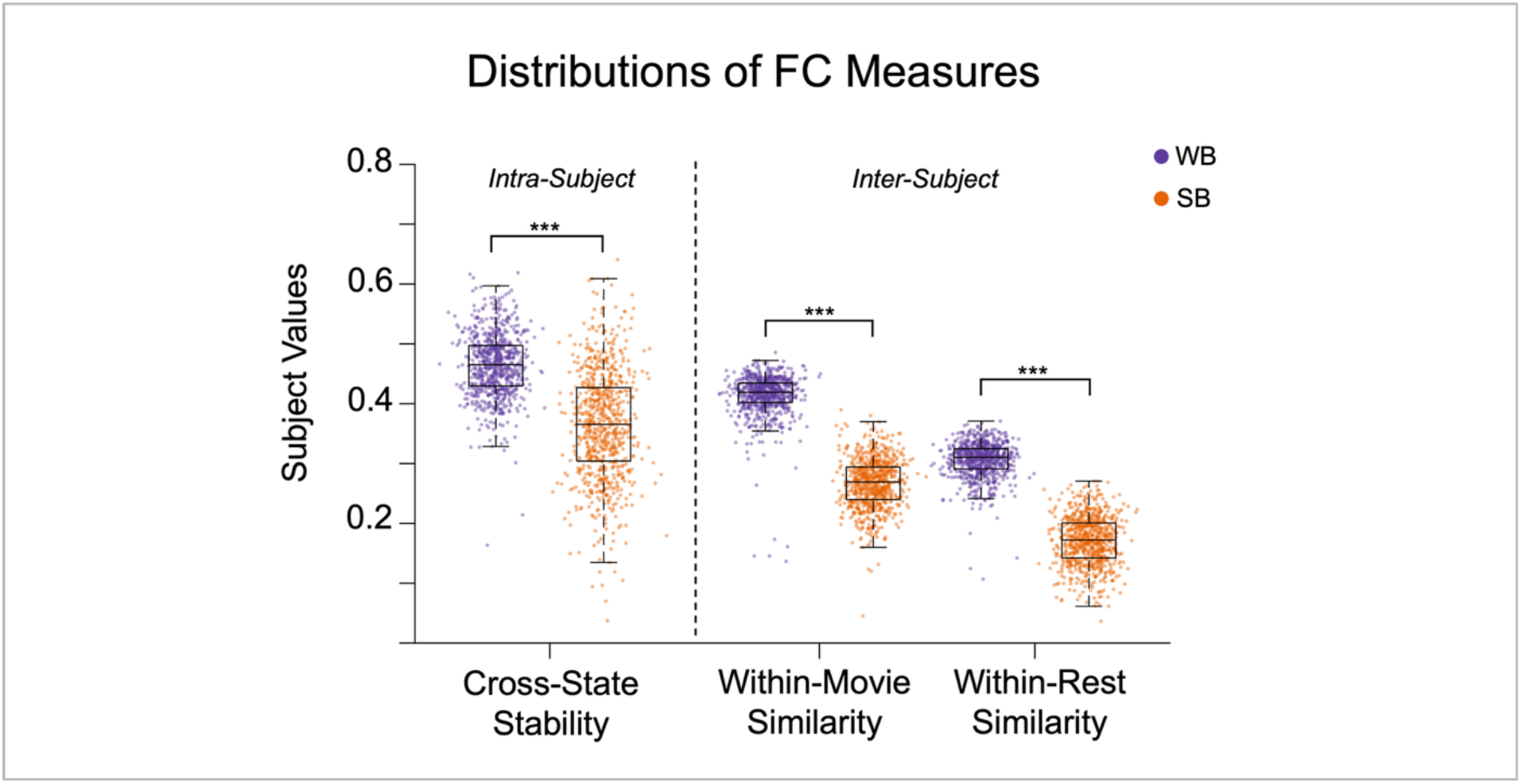
Distributions of FC Measures. Group means and spreads of intra-subject (cross-state stability) and inter-subject (within-state similarity) functional connectivity measures calculated using whole brain (WB) and social brain (SB) connectomes (N=769). Black asterisks denote a statistically significant difference in the mean and dispersion between WB and SB measures (*p*<0.001***).

Using an LME model, higher within-state similarity was positively associated with higher cross-state stability in the whole brain (*β* = 0.265, 95%CI[0.132, 0.398], *p* = 9.61 × 10⁻^05^) and in the social brain (*β* = 1.23, 95%CI[1.12, 1.34], *p* = 2.64 × 10⁻^93^). Furthermore, this relationship was significantly stronger in the social brain relative to the whole brain (*β* = 0.964, 95%CI[0.81, 1.12], *p* = 4.74 × 10⁻^33^). See *Figure 5A*.

**Figure 5.**
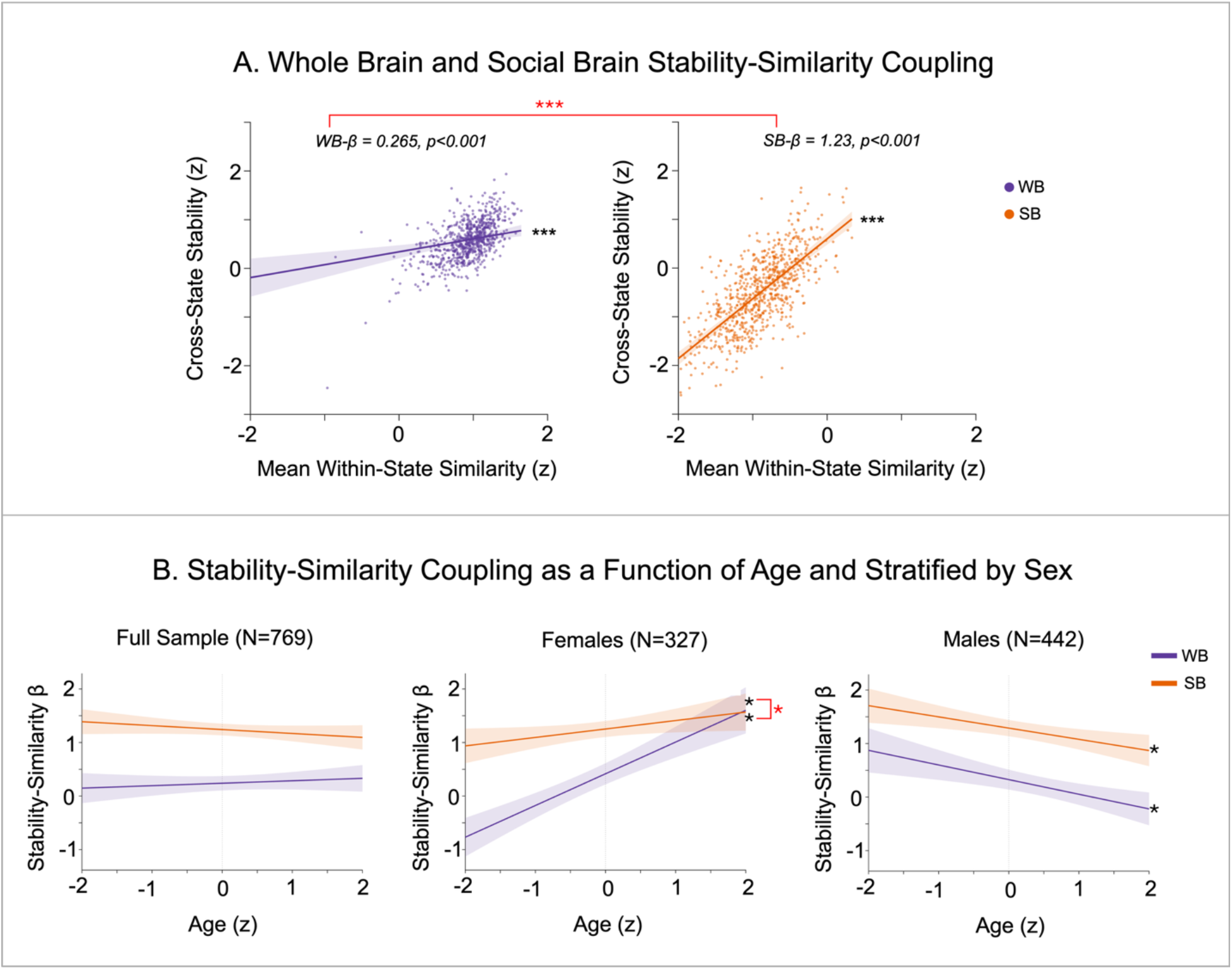
Stability-Similarity Coupling. **(A)** Relationships between cross-state stability and mean within-state similarity estimated for whole brain and social brain (N = 769). Points represent individual subjects’ z-scored values. Solid lines show the model-estimated fixed-effects relationships between stability and similarity for whole brain and social brain, adjusted for age, sex, and mean RMSD, with shaded regions indicating 95% confidence intervals. Black asterisks denote statistically significant slopes for WB and SB separately, and red asterisks indicate a statistically significant difference between WB and SB slopes (\*\*\**p*<0.001). **(B)** Model-estimated stability–similarity coupling as a function of age for the full sample and stratified by sex for whole brain and social brain. Lines show fixed-effect regression slopes (*β*) and shaded regions indicate 95% confidence intervals. Black asterisks denote a significant age moderation effect. Red asterisks denote a significant difference in age moderation between whole brain and social brain.

Across the full sample, age did not appear to moderate the strength of stability-similarity coupling in the whole brain (*β* = 0.046, 95%CI[-0.068, 0.160], *p* = 0.432) or social brain (*β* = - 0.073, 95%CI[-0.174, 0.028], *p* = 0.155). However, when stratifying by sex, males showed a decrease in stability-similarity coupling as a function of age in the whole brain (*β* = -0.273, 95%CI[-0.430, -0.117], *p* < .001) and social brain (*β* = -0.210, 95%CI[-0.342, -0.077], *p* = 0.002), while females showed an increase in stability-similarity coupling as a function of age in the whole brain (*β* = 0.592, 95%CI[0.420, 0.765], *p* < 0.001) and social brain (*β* = 0.158, 95%CI[0.008, 0.308], *p* < 0.05). In females, this age moderation effect was significantly weaker in the social brain relative to the whole brain (*β* = -0.435, 95%CI[-0.637, -0.233], *p* < 0.001). See *Figure 5B* for visualization of the age moderation effects; individual data points are not shown as stability-similarity coupling is estimated across participants.

These findings demonstrate that individuals with more stable connectomes also tend to have FC patterns that are more similar to mature subjects. Although stability and maturational similarity showed more inter-individual variability within the social brain, the strength of their coupling was stronger in the social brain compared to the whole brain. Furthermore, the strength of this stability-similarity coupling was moderated by age in opposite directions for males and females.

### 3.4 Age and SRS Associations with Stability and Maturational Similarity

FC relationships with age and social functioning scores across the full sample and stratified by sex are depicted in *Supplementary Figure 4*. No individual age or SRS effects survived FDR correction (*q* > 0.05). See *Supplementary Tables 1* and *2* for full results.

### 3.5 Multivariate Predictions of Age

Edgewise FC measures for whole brain (35778×1 vectors) and social brain (630×1 vectors) were calculated and used to predict age using regularized ridge regression. See *Supplementary Figure 5*. Whole brain models significantly predicted age from edgewise measures of cross-state stability (mean *r* = 0.400, mean RMSE = 3.22, *p*<0.001), within-rest similarity (mean *r* = 0.550, mean RMSE = 2.97, p<0.001), and within-movie similarity (mean *r* = 0.611, mean RMSE = 2.81, *p*<0.001). Model performance was significantly better for within-movie compared to within-rest similarity predictions of age using r (mean difference = 0.061, 95% CI [0.059, 0.064], *t*(99) = 46.2, *p*<0.001) and RMSE (mean difference = -0.169, 95% CI [-0.174, -0.164], *t*(99) = -66.6, *p*<0.001).

Two of the three social brain models also significantly predicted age: within-rest similarity (mean *r* = 0.158, *p* = 0.002, mean RMSE = 3.53, *p*<0.001) and within-movie similarity (mean *r* = 0.183, *p* = 0.002, mean RMSE = 3.51, *p*<0.001), although with much lower accuracy than whole brain models. Again, model performance was significantly better for within-movie vs. within-rest similarity using *r* (mean difference = 0.026, 95% CI [0.019, 0.031], *t*(99) = 8.86, *p*<0.001) and RMSE (mean difference = -0.027, 95% CI [-0.034, -0.019], *t*(99) = -7.31, *p*<0.001).

### 3.6. Spatial Distribution of Highly Predictive Whole Brain Edges

Predictive performance for whole brain models was higher than social brain models, thus, we only explored the spatial distributions of whole brain edges. Given that previous research suggests the social brain is highly embedded within the larger brain architecture (McCormick et al., 2018; Maliske and Kanske, 2022), we examined the extent to which social brain regions contributed to highly predictive connections in our whole brain models. See *Figure 6*.

**Figure 6.**
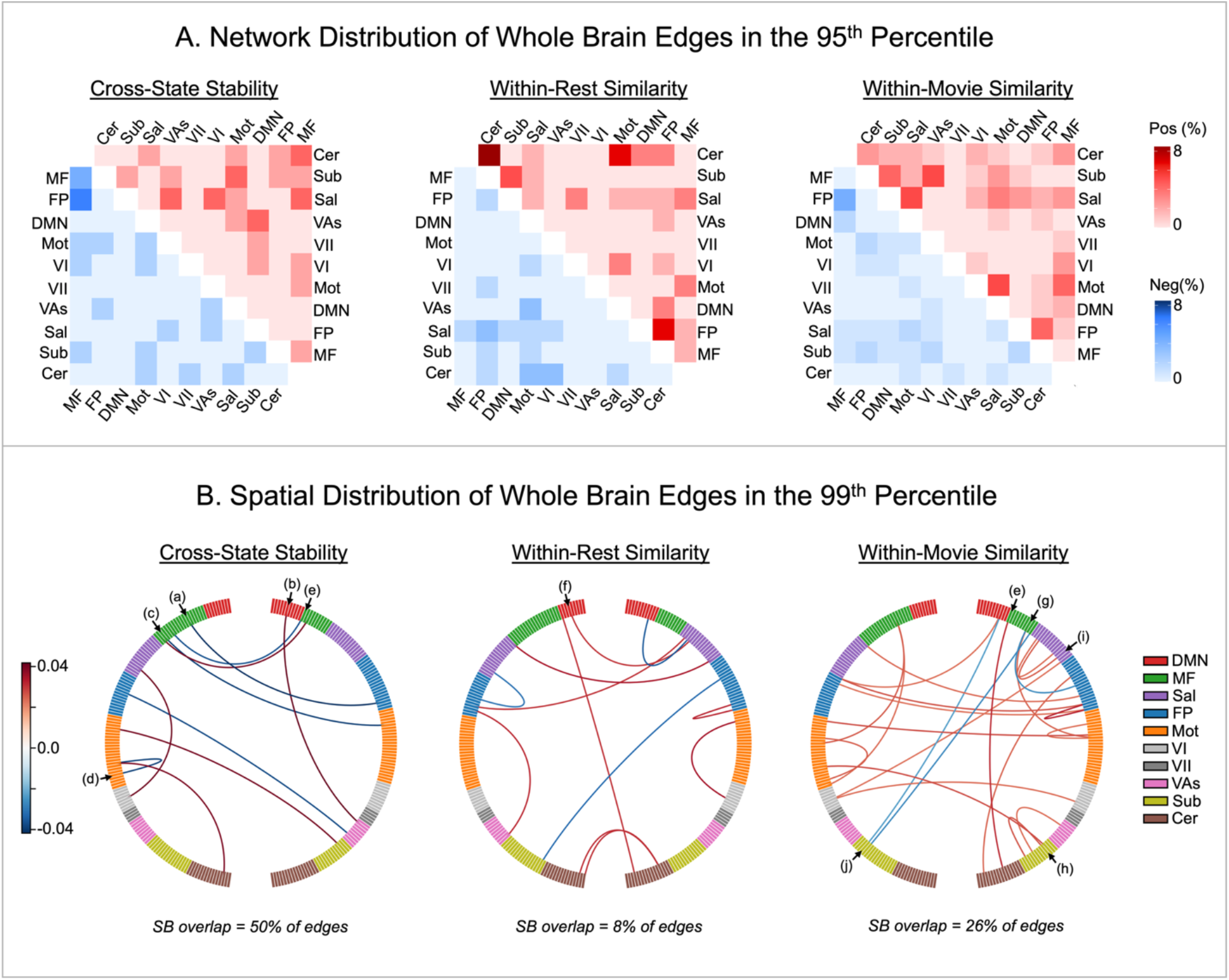
Whole Brain Connections Highly Predictive of Age. **(A)** Heatmaps show the proportion of positive (upper triangle, red) and negative (lower triangle, blue) edges within each network pair, expressed as a percentage of all edges in the 95^th^ percentile. Pos/Neg Edges = total proportion of positive vs negative edges in the 95^th^ percentile across all network pairs. **(B)** Circular network plots visualizing edges in the 99^th^ percentile arranged by hemisphere and network. Line color encodes the sign and magnitude of the contribution weights; warmer colors = positive contributions, cooler colors = negative contributions; greater color intensity reflects larger contribution magnitude. SB overlap = percentage of whole brain edges in the 99^th^ percentile that contain social brain regions. whole brain nodes that correspond to social brain regions are indicated by letter (a = lIFG, b = rTPJ, c = lpSTS, d = lMTG, e = rpSTS, f = pMCC, g = rIFG, h = rNAcc, i = aMCC, j = lHC). Whole brain networks: MF, medial frontal; FP, frontoparietal; DMN, default mode; Mot, Motor; VI, visual I; VII, visual II; VAs, visual association; Sal, salience; Sub, subcortical; Cer, cerebellar.

First, edge contribution weights from each predictive model were estimated using previous methods (e.g. Gao et al., 2019; Greene et al., 2020). For each repetition, fold-specific ridge coefficients were saved, resulting in 1000 ridge coefficients per edge. Given that feature selection was re-run for each fold, only edges that were “reliably selected” across 90% of all model iterations were evaluated. For each selected edge, the mean ridge coefficient (β) was aggregated across folds and repetitions. Edge contributions were then calculated as (β×SD), where SD is the standard deviation of an edge’s value across participants.

Distributions of highly predictive edges were visualized at the network-level (thresholding at the 95^th^ percentile) using signed heatmaps and at the edge-level (thresholding at the 99^th^ percentile) using circular network plots (MNE-Python package: Gramfort et al., 2013). As hypothesized, highly predictive connections were diffusely distributed across the brain and showed both positive and negative edge weights (positive edges = connections where greater stability or similarity predicted higher age, negative edges = connections where lower stability or similarity predicted higher age).

After mapping social brain regions onto the Shen-268 atlas (see *Supplementary Table 3*), we assessed the extent to which highly predictive whole brain connections overlapped with our social brain mask. Overall, there was considerable social brain representation among predictive whole brain edges in the 99^th^ percentile. The proportions of edges containing social brain regions were 50% for cross-state stability, 8% for within-rest similarity, and 26% for within-movie similarity. Many of these social brain regions corresponded to higher (e.g., rTPJ and lMTG) and intermediate-level networks (e.g. aMCC and bilateral IFG), although nodes from visual-sensory (e.g. bilateral pSTS) and limbic networks (e.g. rNAcc, lHC) were also evident. Interestingly, across all models, highly predictive connections that overlapped with social brain regions all interfaced with non-social regions, indicating that inter-network connectivity involving the social brain may better capture multivariate developmental effects relative to intra-network connectivity within the social brain.

## 4. Discussion

The goal of this study was to assess inter- and intra-subject measures of FC in children and adolescents, with a focus on the social brain. Using a large sample of fMRI data from the HBN Biobank, we computed the similarity of each participant’s FC matrix to a mature reference group. We also computed cross-state stability to capture how much modulation occurred for each participant between movie and rest. Linear mixed effects models tested for age-based effects, and we conducted stratified analyses to test for sex-specific effects. Overall, we showed that cross-state stability and similarity to mature peers are robustly coupled such that individuals with more stable FC across states tend to exhibit more mature FC architecture. We also newly demonstrated that this coupling is even stronger in the social brain and shows age-related variation in sex-specific ways. When assessing each FC measure independently, we found small univariate relationships with age and social functioning scores that did not survive FDR correction. In contrast, our multivariate predictions of age performed much better, demonstrating that retaining edgewise information better captures complex and spatially heterogenous developmental effects that are lost when collapsing across edges with single summary measures.

### 4.1 Whole brain and social brain connectomes are discriminable in youth

Using a binary classification approach called fingerprinting (Finn & Shen et al., 2015), whole brain FC patterns have been characterized as unique and identifiable in adults (Finn & Shen et al., 2015; Vanderwal et al., 2017), as well as in children and adolescents (Jalbrzikowski et al., 2020; Horien et al., 2019; Demeter et al., 2020; Camp et al., 2024). Consistent with these findings, we found that pediatric whole brain FC patterns show high identifiability and discriminability, particularly within higher-level association networks (as shown in Finn & Shen et al., 2015; Horien et al., 2019). We also provide novel evidence that social brain FC is individualized in youth ages 6-21, and that this uniqueness is best captured using continuous measures of discriminability, compared to binary fingerprinting measures.

The social brain has considerable overlap with the default mode network (Mars et al., 2012), and these regions show substantial functional reconfiguration during movie-watching (Yeshurun et al., 2021; Redcay and Moraczewski, 2020). Given that we assessed fingerprinting across rest and movies, we argue that lower social brain cross-state identifiability may reflect greater state-dependent reorganization rather than weaker individualization per se. In other words, it is possible that a fingerprinting algorithm using movie-movie or rest-rest pairings might show different results in this regard.

### 4.2 Stability-similarity coupling is enhanced in the social brain and is moderated by age in sex-dependent ways

The stability of FC across time and tasks is a well-known feature of the mature connectome (Gratton et al., 2018). At the same time, mature FC patterns also contain a canonical network organization that is broadly similar across individuals (Yeo et al., 2011). Although these features have typically been characterized independently, recent work has demonstrated that there are robust positive relationships between FC stability and group-similarity (Vanderwal et al., 2021; Liao et al., 2021; Corriveau et al., 2023). Building on this work, we newly demonstrate that more stable FC patterns in children and youth also exhibit FC patterns that are more similar to mature architecture, and that this stability-similarity coupling is even stronger in the social brain. This is intriguing given that cross-state stability showed greater variability across subjects in the social brain. These findings are consistent with the notion that default mode regions of the social brain show idiosyncratic responses to movies (Yeshurun et al., 2021; Redcay and Moraczewski, 2020). Although this sensitivity to movies appears to decrease cross-state identifiability in the social brain, it simultaneously also appears to capture individual variability related to the strength of stability-similarity coupling.

Our findings also suggest that the developmental organization of this stability–similarity coupling differs by sex. Specifically, we found that stability-similarity coupling increased with age in females but decreased with age in males. Prior work has demonstrated that during development, females typically show greater network segregation, while males show greater between-network integration (Satterthwaite et al., 2015; Wu et al., 2013). Here, our results indicate that sex-dependent developmental differences may not only be reflected in the strength of individual connections, but also in how different organizational properties of FC are coupled.

In females, we also found that the positive relationship between age and stability-similarity coupling was weaker in the social brain relative to the whole brain. This smaller age-related variation in the social brain raises an intriguing possibility that the social brain may already be relatively mature in females by early childhood. This notion is consistent with several behavioural studies demonstrating that during youth, females tend to have more advanced social cognitive abilities compared to males of the same age (e.g. Calero et al., 2013; Poznyak et al., 2019; Desatnik et al., 2023).

### 4.3 Summary FC measures do not relate to age or social functioning scores

Prior work using ISCs has shown that similarity of movie-evoked neural responses relative to mature subjects increases with age and social cognitive abilities in higher-level social brain regions (Richardson et al., 2018, 2019; Moraczewski et al., 2018, 2020). In contrast to these findings and our hypotheses, we did not find significant positive associations between maturational similarity and age or social functioning scores, nor did we find stronger effects in the social brain or with movie-watching versus rest. Furthermore, while we anticipated cross-state stability to be positively related to age and social functioning scores, our results did not demonstrate this. Interestingly, these findings are at odds with our previous work in a smaller sample size (N=210) in which social functioning scores showed positive relationships with cross-rest stability and within-movie group similarity across the whole brain (Vanderwal et al., 2021). When using a much larger sample size (N=769), these results did not replicate. However, it is worth noting that the FC measures used here differed in that we specifically queried rest-to-movie stability and similarity relative to mature subjects rather than the group as a whole.

Although we found that stability-similarity coupling varies with age in sex-specific ways, none of the age or SRS associations with individual FC measures survived FDR correction in our sex-stratified models. Nonetheless, some opposing developmental patters were evident between males and females (see *Supplemental Figure 4* and *Supplementary Tables 1* and *2*). However, because sex-stratified models only test for associations within each sex separately, it is possible that testing for sex-by-age interactions across the full sample may reveal sex-based differences in these effects. In healthy adults, sex-by-age interactions in FC during movie-watching were recently reported, with the strongest effects observed in visual regions (Yang et al., 2025). This study also showed an effect of movie clip type on sex and age relationships, particularly in the limbic network. We could not assess the effect of movie type here, but it will be an important consideration in future developmental work.

### 4.4 Edgewise stability and similarity measures capture multivariate developmental patterns

Developmental effects that may be spatially heterogeneous across different connections may be missed with summary measures that do not preserve information about which connections show stronger or weaker stability and similarity as a function of age. Thus, to preserve edgewise information, we characterized each participant’s stability and similarity using vectors where elements represent connection-level values. Using multivariate predictive modelling (e.g., Gao et al., 2019), we found that edgewise stability and maturational similarity measures captured stronger whole brain developmental relationships compared to our univariate summary measures. Consistent with our hypotheses, age-based effects were spatially heterogeneous across the brain, as some connections became more stable or similar to mature subjects with age, whereas others became increasingly differentiated. These divergent patterns suggest that maturation of FC architecture does not follow a single uniform trajectory across the brain and spatially sensitive methods are needed to identify these developmental patterns.

Although our social brain models showed lower performance, many of the highly predictive edges in our whole brain models contained social brain regions. Importantly, these highly predictive social brain connections all interfaced with regions outside the social brain. Thus, the lower performance of the social brain models may reflect the fact that our social brain mask excluded connections to the rest of the brain. Together, these findings highlight the importance of studying the social brain as an embedded system within larger brain circuitry rather than as an isolated functional unit (McCormick et al., 2018; Maliske and Kanske, 2022).

### 4.5 Maturational similarity during movies predicts age more accurately than during rest

Movies have been shown to draw out individual variability not otherwise detectable with traditional task (Cantlon and Li, 2013; Richardson et al., 2019) or resting-state paradigms (Finn and Bandettini, 2021). Here, our multivariate findings further advocate the use of movies for studying development as we observed better predictive performance of age in our maturational similarity models during movie-watching compared to rest, for both whole brain and social brain. The fact that movies better capture individual variability is intriguing given that movies synchronize neural responses across many regions of the brain (Hasson et al., 2004). The proposed interpretation of this paradoxical finding, which we again put forth here, is that by reducing overall between-subject variability, meaningful individual variability is more easily identified (Vanderwal et al., 2017; Finn et al., 2017). These findings, along with others (e.g. Redcay and Moraczewski, 2020), highlight the advantage of using naturalistic paradigms containing dynamic and contextually embedded social stimuli for capturing individual differences and developmental variability.

### 4.6 Strengths and Limitations

A main limitation of the current study is that the data are from a psychiatrically enriched sample, and we do not yet know the nature of the studied relationships in a typically developing population. In other words, the HBN data are a convenience sample, used here because it is the only developmental dataset of this size to include movie-fMRI. It is thus possible that results (especially age-related effects) are confounded by the inclusion of multiple neurodevelopmental disorders and clinical heterogeneity of varying severities across different ages. On the other hand, while studying functional organization in typically developing children is needed to characterize normative patterns of brain maturation, such findings may not be representative of real-world populations, as approximately 30% of youth in Western countries are estimated to have a psychiatric diagnosis (Tatsiopoulou et al., 2025). By including sex-stratified analyses, we also provide insight into whether social brain development in a psychiatrically enriched sample follows similar or distinct patterns in males and females. Given that social functioning (Proverbio, 2023) and psychiatric vulnerabilities differ between males and females (Wiglesworth et al., 2023; López-Ojeda & Hurley, 2021; Christiansen et al., 2022), examining how social brain development differs by sex may help clarify the role of the social brain in these sex-specific developmental risks. Unfortunately, because measures related to gender identity were not available, we were not able to disentangle sex-based and gender-based effects in the current study. Especially in the studied age range, this should be included where possible in future work.

There are several additional limitations worth noting. First and foremost, the data are cross-sectional and this precludes inferences about developmental change. Future longitudinal work is needed to determine whether these observed age-related patterns reflect individual developmental trajectories. Second, because reliable estimates of FC require sufficient scan lengths, we did not assess developmental patterns related to within-state FC stability (i.e., rest– rest or movie–movie). Our findings are thus confined to stability assessed across rest and movie states. Future studies are needed to assess stability both within- and across various brain states. Finally, while we assessed maturational similarity relative to the healthiest, oldest participants available in the sample (ages 17–21 years) as an index of mature connectomes, future work should aim to assess maturational similarity relative to adults who are further along in prefrontal cortical development (Sydnor et al., 2021). This may provide a more accurate benchmark of mature functional organization that would amplify or change the findings reported here.

## 5. Conclusions

The current study provides novel evidence that rest-to-movie stability within a subject is strongly related to peer-to-peer similarity, such that individuals with more stable FC across states tend to have FC patterns that are more similar to mature peers. Furthermore, we newly show that this stability-similarity coupling is enhanced in the social brain and that age moderates the strength of this relationship in a sex-dependent manner. Univariate effects relating each FC measure independently to age and social functioning were small and nominally significant. In contrast, our multivariate predictions of age performed much better. Thus, preserving edgewise information about which connections show stronger or weaker stability and similarity with age captures spatially heterogeneous effects that are missed when collapsing across edges. In our whole brain models, many highly predictive edges overlapped with our social brain mask, however, these social brain regions were always connected to regions outside the social brain. This later finding highlights the importance of studying the development of social brain systems embedded within the larger whole brain architecture.

