## Supplemental Figures and Tables for "Cross-state stability and maturational similarity of the social brain connectome in children and adolescents"

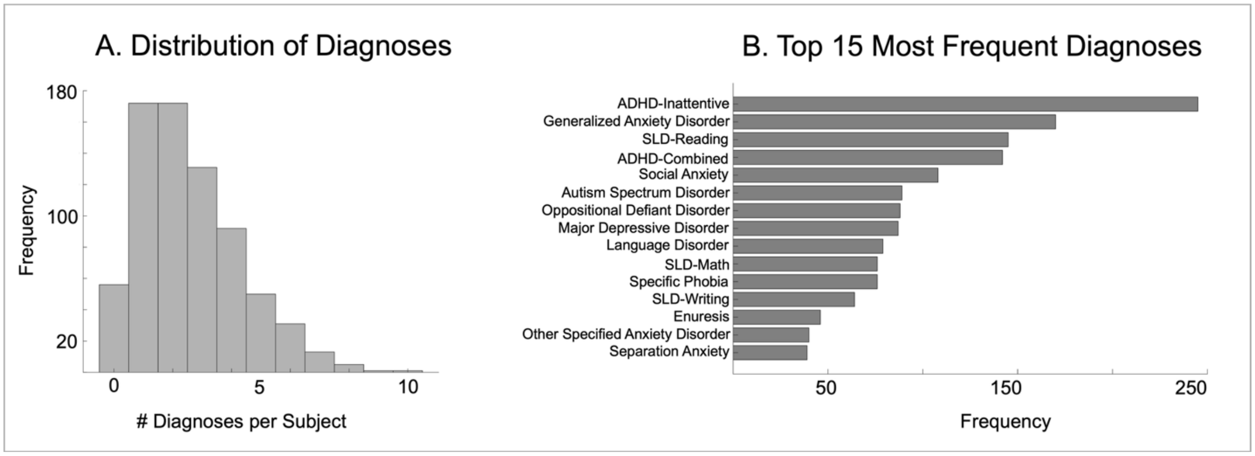


**Supplemental Figure 1. Psychiatric Diagnoses for 724 participants.**
**(A)** Distribution of the number of clinician consensus DSM-5 psychiatric diagnoses per subject among participants who completed the semi-structured diagnostic interviews (N = 724).
**(B)** Frequencies of the 15 most common diagnoses within this subsample of participants with complete diagnostic data (N = 724); ADHD = Attention-Deficit/Hyperactivity Disorder; SLD = Specific Learning Disorder.

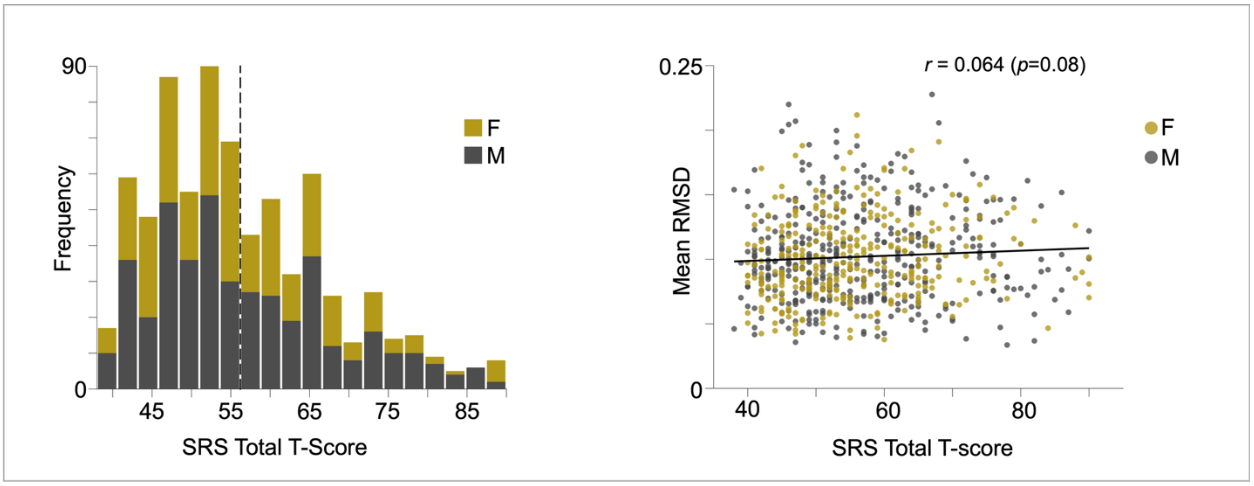


**Supplementary Figure 2.** Distributions for SRS scores (N=736, female = 314) are visualized by sex with group means indicated by the dashed lines (left). The relationship between head motion (mean RMSD) and SRS scores are visualized by sex (right).

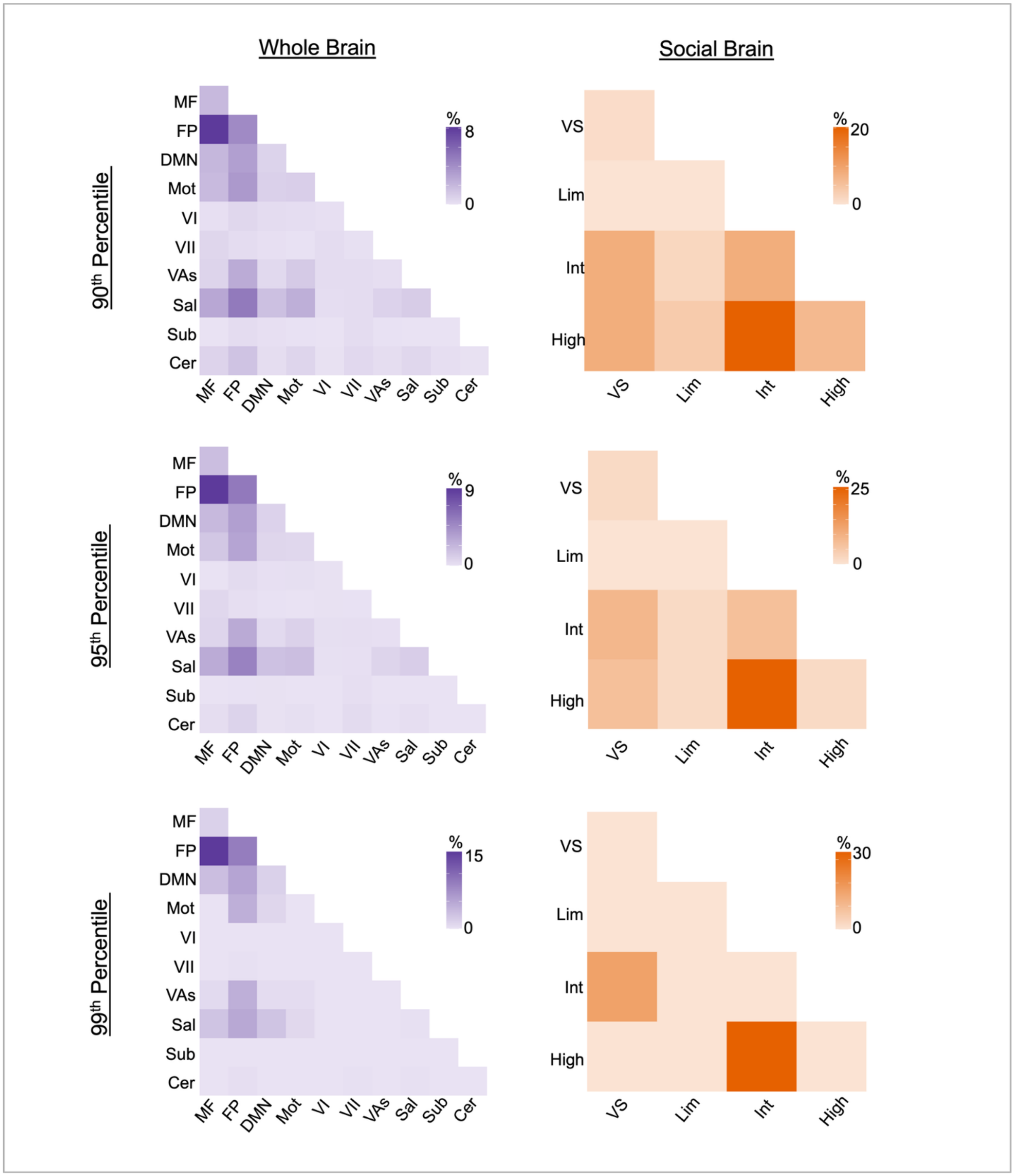


**Supplementary Figure 3.** Network-level distribution of high differential power (DP) edges, thresholding at 90th, 95th, and 99th percentiles. Tile colour intensity indicates the proportion of high-DP edges assigned to each network pair. Whole brain networks: MF, medial frontal; FP, frontoparietal; DMN, default mode; Mot, Motor; VI, visual I; VII, visual II; VAs, visual association; Sal, salience; Sub, subcortical; Cer, cerebellar. Social brain networks: VS, visual-sensory; Lim, limbic; Int, intermediate; High, higher-level.

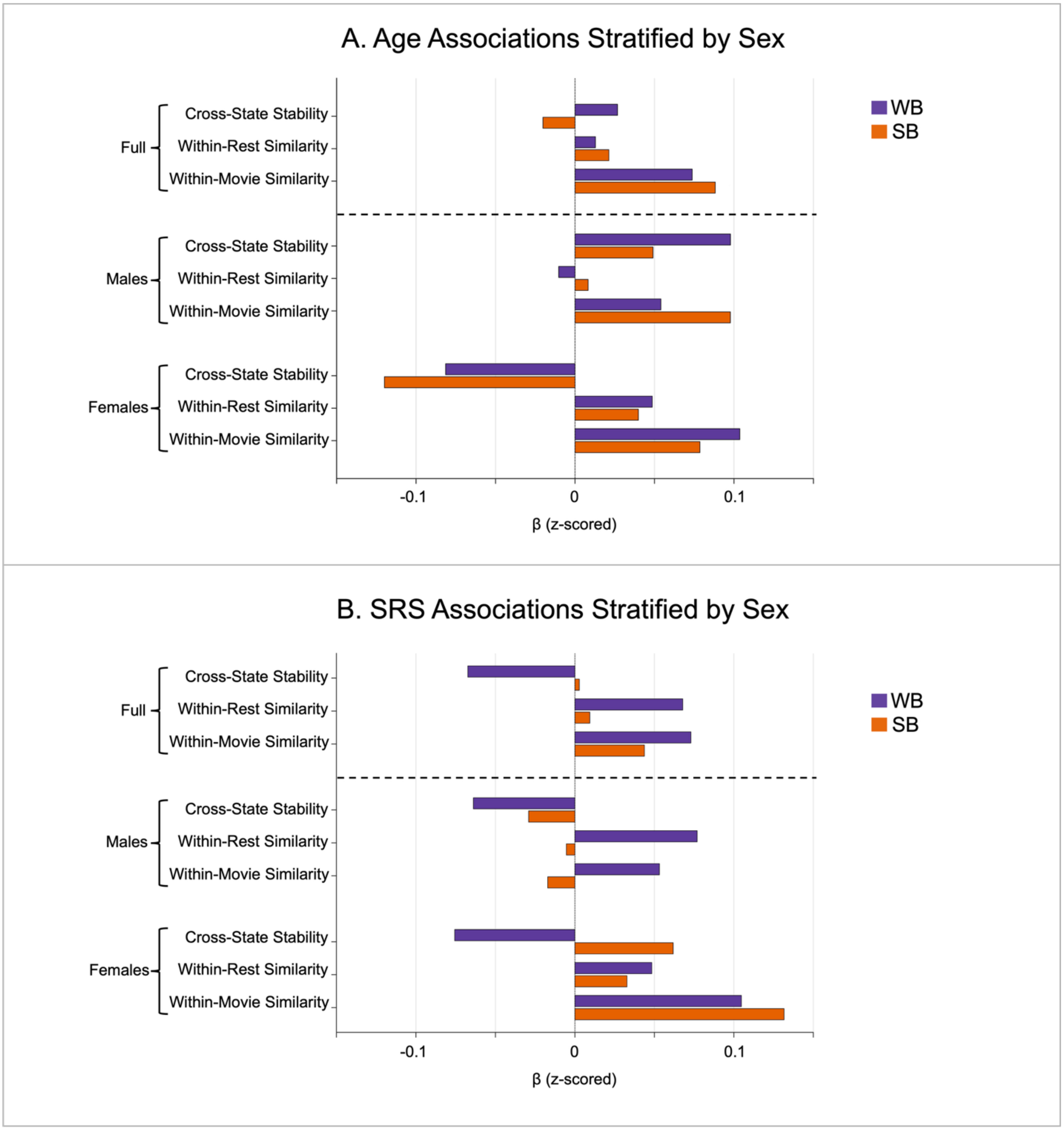


**Supplementary Figure 4. Univariate associations for each FC measure**
Standardized $\beta$ coefficients explain variability in each whole brain (WB) and social brain (SB) functional connectivity (FC) metric from **(A)** Age across the full sample (N=769) and stratified by sex (males=442; females=327), and **(B)** Social Responsiveness Scale-2 (SRS) total T-scores across the full sample (N=736) and stratified by sex (males=422; females=314). The direction of SRS scores are flipped so that higher scores indicate better functioning. No effects remained significant after FDR-correction ($q$ < .05).

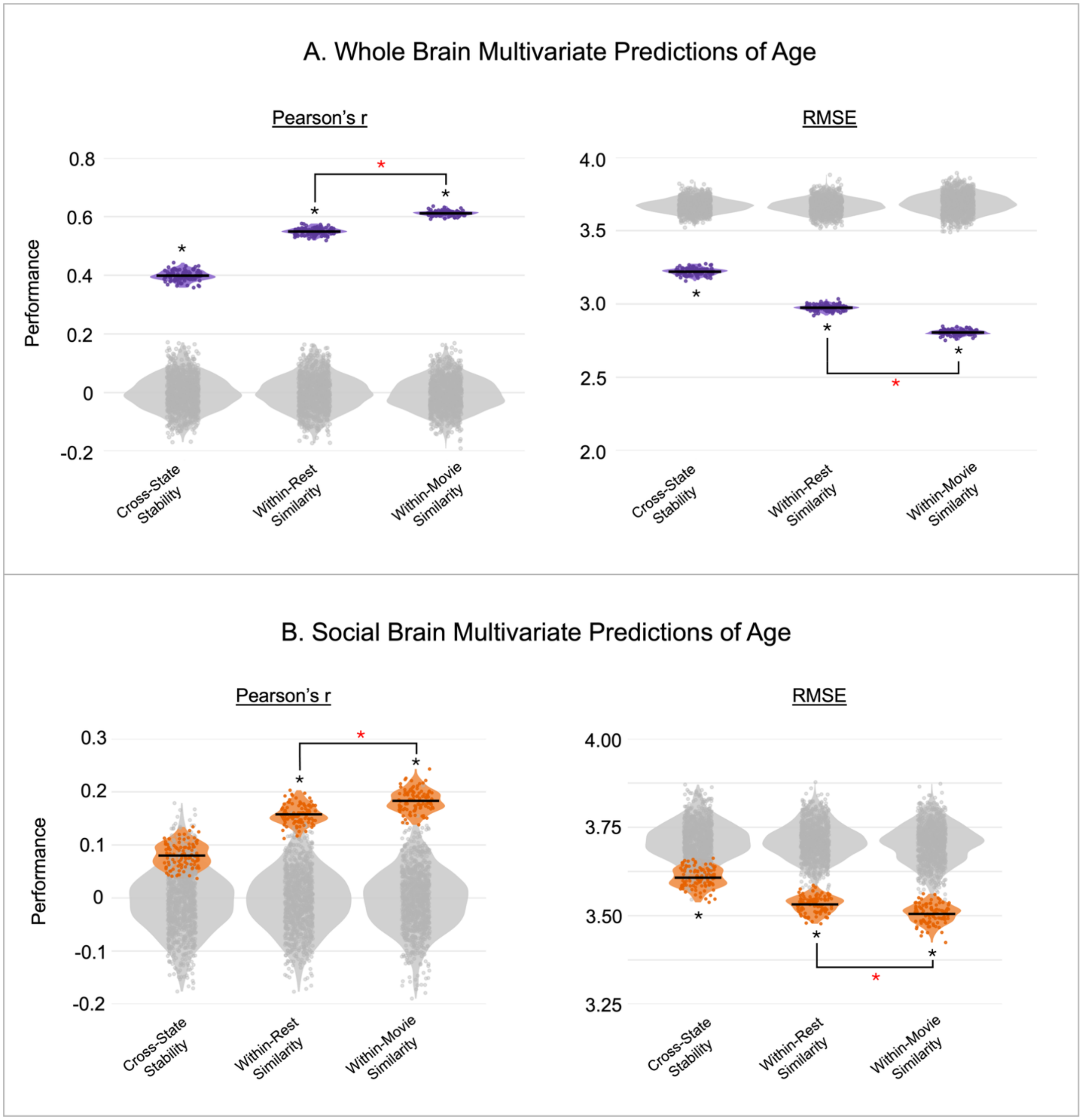


**Supplementary Figure 5. Multivariate Predictions of Age using Ridge Regression**
**(A)** Whole Brain and **(B)** Social Brain multivariate predictions of age were assessed with Pearson correlation (r; left panel) and root-mean squared error (RMSE; right panel). Performance metrics from 100 repeated model fits are shown in purple for whole brain and orange for social brain. Performance distributions from 1000 null models are shown in grey. Black stars denote statistically significant predictive performance of each model. Red stars denote statistically significant difference in model performance for Within-Movie Similarity vs Within-Rest Similarity.

| FC Measure | Effect Type | Full Sample (N=769) β [95% CI] | | Males (N=442) β [95% CI] | | Females (N=327) β [95% CI] | |
|---|---|---|---|---|---|---|---|
| Cross-State Stability | WB | 0.027 [−0.043, 0.096] | n.s. | 0.098 [0.005, 0.190] | * | −0.081 [−0.185, 0.022] | n.s. |
| | SB | −0.020[−0.089, 0.049] | n.s. | 0.049 [−0.043, 0.141] | n.s. | −0.120 [−0.223, −0.017] | * |
| | SB - WB | -0.047 [-0.120, 0.026] | n.s. | -0.049 [-0.145, 0.048] | n.s. | -0.039 [-0.152, 0.075] | n.s. |
| Within-Rest Similarity | WB | 0.013 [−0.056, 0.082] | n.s. | −0.010 [−0.102, 0.082] | n.s. | 0.049 [−0.056, 0.153] | n.s. |
| | SB | 0.021 [−0.048, 0.090] | n.s. | 0.008 [−0.083, 0.100] | n.s. | 0.040 [−0.064, 0.144] | n.s. |
| | SB - WB | 0.008 [-0.061, 0.078] | n.s. | 0.019 [-0.072, 0.109] | n.s. | -0.009 [-0.119, 0.101] | n.s. |
| Within-Movie Similarity | WB | 0.074 [0.002, 0.145] | * | 0.054 [−0.042, 0.150] | n.s. | 0.104 [−0.005, 0.212] | n.s. |
| | SB | 0.088 [0.017, 0.160] | * | 0.098 [0.002, 0.193] | * | 0.079 [−0.030, 0.187] | n.s. |
| | SB - WB | 0.015 [-0.056, 0.085] | n.s. | 0.044 [-0.052, 0.139] | n.s. | -0.025 [-0.128, 0.078] | n.s. |

**Supplementary Table 1.** Results of linear mixed-effects models examining associations between age and whole brain (WB) and social brain (SB) functional connectivity (FC) measures in the full sample and stratified by sex. Values represent standardized β coefficients and 95% confidence intervals. SB–WB represents the Age x Connectome Type (WB/SB) interaction effect, calculated as $\beta$SB − $\beta$WB. * indicates nominal significance ($p$ < .05) that did not survive FDR correction; n.s. indicates $p \geq .05$.

| FC Measure | Effect Type | Full Sample (N=736) β [95% CI] | | Males (N=422) β [95% CI] | | Females (N=314) β [95% CI] | |
|---|---|---|---|---|---|---|---|
| Cross-State Stability | WB | -0.067 [-0.133, -0.002] | * | -0.064 [-0.148, 0.021] | n.s. | -0.076 [-0.180, 0.029] | n.s. |
| | SB | 0.003 [-0.063, 0.069] | n.s. | -0.029 [-0.114, 0.055] | n.s. | 0.062 [-0.043, 0.167] | n.s. |
| | SB - WB | 0.070 [-0.003, 0.143] | n.s. | 0.035 [-0.058, 0.128] | n.s. | 0.137 [0.019, 0.256] | * |
| Within-Rest Similarity | WB | 0.068 [0.002, 0.133] | * | 0.077 [-0.007, 0.161] | n.s. | 0.048 [-0.057, 0.154] | n.s. |
| | SB | 0.009 [-0.056, 0.075] | n.s. | -0.005 [-0.089, 0.078] | n.s. | 0.033 [-0.073, 0.138] | n.s. |
| | SB - WB | -0.058 [-0.129, 0.012] | n.s. | -0.082 [-0.171, 0.006] | n.s. | -0.016 [-0.133, 0.102] | n.s. |
| Within-Movie Similarity | WB | 0.073 [0.005, 0.141] | * | 0.053 [-0.035, 0.141] | n.s. | 0.105 [-0.004, 0.213] | n.s. |
| | SB | 0.044 [-0.024, 0.112] | n.s. | -0.017 [-0.105, 0.071] | n.s. | 0.132 [0.023, 0.240] | * |
| | SB - WB | -0.029 [-0.099, 0.041] | n.s. | -0.070 [-0.161, 0.020] | n.s. | 0.027 [-0.085, 0.138] | n.s. |

**Supplementary Table 2.** Results of linear mixed-effects models examining associations between SRS scores and whole-brain (WB) and social-brain (SB) functional connectivity (FC) measures in the full sample and stratified by sex. Values represent standardized β coefficients and 95% confidence intervals. SB–WB represents the SRS x Connectome Type (WB/SB) interaction effect, calculated as $\beta$SB − $\beta$WB. Scores for SRS were reversed so that higher values indicate better functioning. * indicates nominal significance ($p$ < .05) that did not survive FDR correction. n.s. indicates $p \geq .05$.

| Social Brain ROI | Region Name | Network | Top Shen Parcel | Coverage (%) |
|---|---|---|---|---|
| aMCC | Anterior mid-cingulate cortex | Intermediate | 15 | 50.00 |
| dmPFC | Dorsomedial prefrontal cortex | High | 145 | 94.50 |
| frontalpole* | Medial frontal pole | High | 140 | 76.00 |
| lAI | Left anterior insula | Intermediate | 155 | 84.50 |
| lamygdala | Left amygdala | Limbic | 228 | 76.50 |
| lCb | Left cerebellum | Intermediate | 253 | 100.00 |
| lFFA | Left fusiform gyrus | Visual-Sensory | 200 | 70.50 |
| lHC | Left hippocampus | Limbic | 231 | 96.00 |
| lIFG | Left inferior frontal gyrus | Intermediate | 151 | 96.00 |
| lIPL | Left inferior parietal lobe | Intermediate | 179 | 64.00 |
| lMTG | Left middle temporal gyrus | High | 197 | 87.50 |
| lMTV5 | Left middle temporal V5 area | Visual-Sensory | 209 | 88.00 |
| lNAcc | Left nucleus accumbens | Limbic | 259 | 100.00 |
| lpSTS | Left posterior superior temporal sulcus | Visual-Sensory | 192 | 77.00 |
| lSMA | Left supplementary motor area | Intermediate | 149 | 75.00 |
| ltemporalpole | Left temporal pole | High | 187 | 84.00 |
| lTPJ | Left temporo-parietal junction | High | 183 | 88.50 |
| PCC | Posterior cingulate cortex | High | 222 | 83.00 |
| PCu | Precuneus | High | 225 | 85.00 |
| pMCC | Posterior mid-cingulate cortex | High | 223 | 81.00 |
| rACC* | Rostral anterior cingulate cortex | Limbic | 140 | 68.00 |
| rAI | Right anterior insula | Intermediate | 36 | 56.50 |
| ramygdala** | Right amygdala | Limbic | 99 | 91.50 |
| rCb | Right cerebellum | Intermediate | 114 | 100.00 |
| rFFA | Right fusiform gyrus | Visual-Sensory | 66 | 60.00 |
| rHC** | Right hippocampus | Limbic | 99 | 37.50 |
| rIFG | Right inferior frontal gyrus | Intermediate | 16 | 88.00 |
| rIPL | Right inferior parietal lobe | Intermediate | 45 | 96.50 |
| rMTG | Right middle temporal gyrus | High | 64 | 96.50 |
| rMTV5 | Right middle temporal V5 area | Visual-Sensory | 74 | 46.50 |
| rNAcc | Right nucleus accumbens | Limbic | 125 | 92.50 |
| rpSTS | Right posterior superior temporal sulcus | Visual-Sensory | 65 | 49.00 |
| rSMA | Right supplementary motor area | Intermediate | 31 | 56.50 |
| rtemporalpole | Right temporal pole | High | 53 | 88.50 |
| rTPJ | Right temporo-parietal junction | High | 50 | 70.00 |
| vmPFC | Ventromedial prefrontal cortex | Limbic | 3 | 46.00 |

**Supplementary Table 3.** For each 36 social brain ROIs, the primary corresponding parcel ("Top Shen Parcel") and ROI coverage (% of ROI voxels contained in the parcel) are reported. In two cases, the best matching Shen parcel was assigned to two different social brain ROIs (these ROIs are denoted by asterisks). To obtain the Top Shen Parcels, the 36 social brain ROIs were mapped onto the Shen-268 atlas using a voxel-based intersection approach. Prior to overlap estimation, both the Shen-268 atlas and all social brain ROI masks were resampled into a common MNI152

1 mm space. For each social ROI, the Shen atlas was masked by the ROI, and the number of overlapping voxels for each Shen parcel was computed. For each social brain ROI, we computed voxel-wise coverage as the percentage of the social brain voxels occupied by the parcel. The Shen parcel exhibiting the greatest number of overlapping voxels was designated as the primary corresponding parcel. Because the social brain ROIs were substantially smaller in spatial extent (200 voxels each) than the Shen-268 parcels (several thousand voxels), the mapping procedure was designed to provide a best-approximation rather than a one-to-one anatomical equivalence.
